# Can a nick promote partial genome re-replication?

**DOI:** 10.1101/2020.11.14.381277

**Authors:** Erik Johansson, John F.X Diffley

**Affiliations:** Chromosome Replication Laboratory The Francis Crick Institute, 1 Midland Road, London, NW1 1AT; Dept. of Medical Biochemistry and Biophysics, Umeå University, 90187 Umeå, Sweden

## Abstract

Single-stranded DNA breaks, including simple nicks, are amongst the most common forms of DNA damage in cells. They can be readily repaired by ligation; however, if a nick occurs just ahead of an approaching replisome, the outcome is a ‘collapsed’ replication fork in which the nick is converted into a single-ended double-strand DNA break. Attention has largely focused on the processes by which this broken end is used to prime replication restart. We realized that in eukaryotic cells, where replication initiates from multiple replication origins, a second fork converging on the collapsed fork offers additional opportunities for repair, but also generates a substrate that can promote localized re-replication. We have modelled this with purified proteins *in vitro* and have found that there is, indeed, an additional hazard that eukaryotic replisomes face. We discuss how this problem might be mitigated.

---

After replication into a nick, one ‘prong’ of the replication fork is converted into a single-ended double strand break whilst a nick is retained on the other prong of the fork (Fig. 1A i and ii). In replication restart (Fig. 1A v), this nick is ligated, the broken end is resected by a 5’->3’ nuclease and a recombinase promotes strand invasion of the single stranded 3’ overhang [1, 2]. A new replisome can then assemble at this primer to complete replication. In *Escherichia coli*, each of the two replication forks from the single chromosomal replication origin (*oriC*) must travel over 2Mb and are terminated in a zone ~180° from the origin; consequently replication restart is the only way to complete replication after fork collapse [1, 3]. In eukaryotes, the genome is replicated from multiple origins and individual replication forks travel much shorter distances than in *E.coli*. There are many ‘dormant’ replication origins which can be activated if needed and, outside ribosomal DNA, there are no programmed termination zones [4]. In contrast to *E.coli*, therefore, a collapsed replication fork will likely be met by a converging replication fork from a downstream origin (Fig. 1A iii). There are three possible outcomes of such an encounter. If the nick generated during the collapse event remains unligated (Fig. 1A iv), the converging fork will also collapse upon reaching the nick, generating the equivalent of a ‘clean’ double strand break, which can be repaired by non-homologous end joining (NHEJ) or homologous recombination (HR) using the sister chromatid as a template [2]. If, however, the nascent DNA is ligated to the nicked template before the converging fork reaches it — regardless of whether the original nick was on the leading or lagging strand template — either replication restart can allow resumption of replication as described above (Fig. 1A v), or the converging fork can continue beyond the position of the nick using the previously replicated DNA as template (Fig. 1A vi). This rogue replication fork will now be following the partner of the original collapsed fork (Fig. 1A vii), potentially re-replicating large regions of the chromosome.

**Figure 1.**
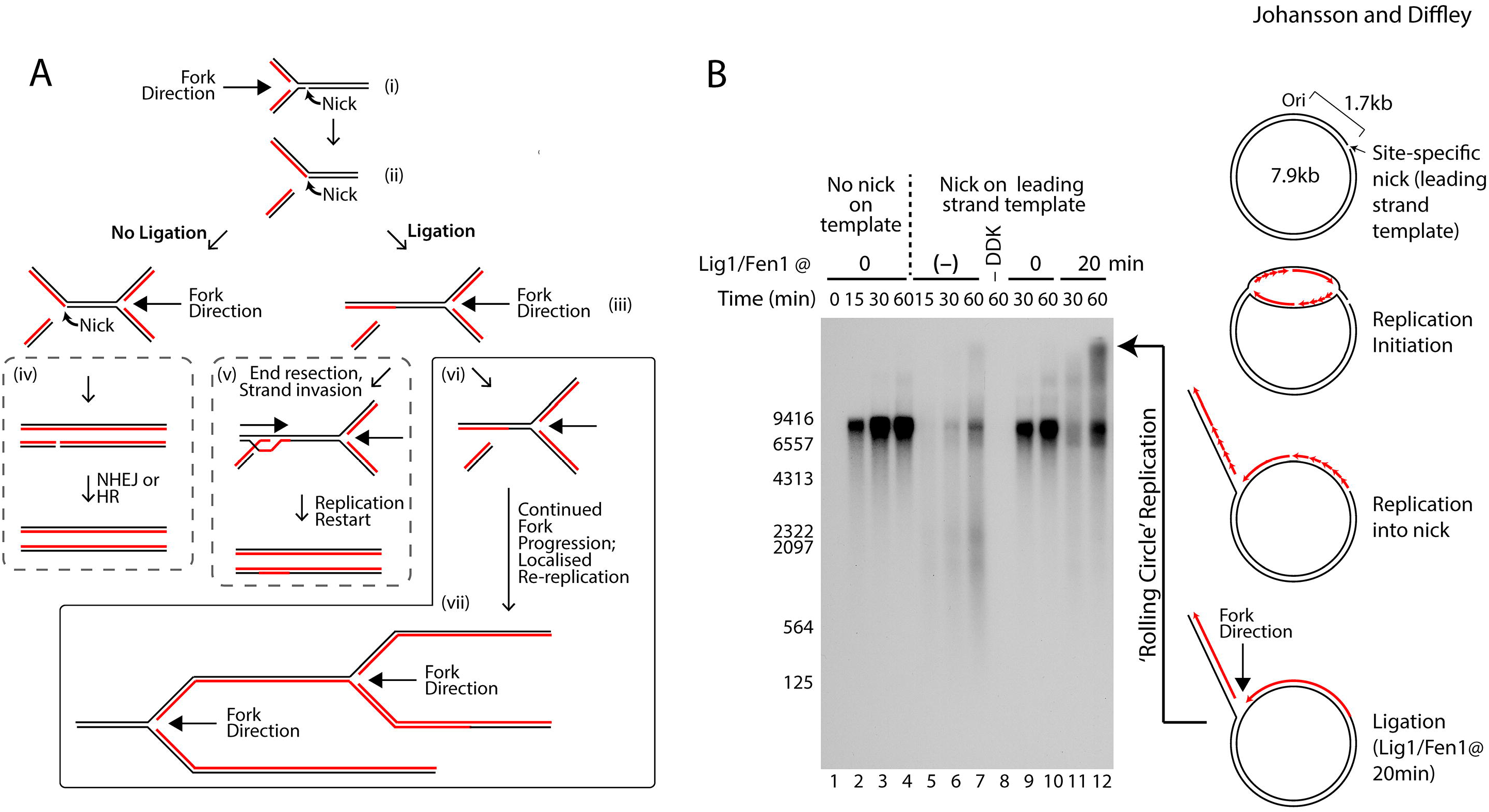
A) Possible outcomes when replication forks encounter a nick on either one of the two templae strands. (i-iii) A nick in a template strand can either remain unligated or be ligated. (iv) In case a nick remains unligated can a converging replication fork rescue the collapsed fork, followed by double-strand break repair by Non-homologous end-joining (NHEJ) or homologous recombination (HR) pathways. (v) Nick ligation between a newly synthesized strand and a template strand allows the resected single-stranded 3′-end to form a D-loop. Leading and lagging strand synthesis is re-established (called homologous recombination-dependent DNA replication (RDR) in *E.coli*; Break induced replication (BIR) in eukaryotes) and is resolved when colliding with the converging fork. (vi) Nick ligation between a newly synthesized strand and a template strand allows the converging fork to continue past the nick site, resulting in re-replication B) Adding Fen1 and ligase after at least one of the two replication forks have met at the nick results in rolling circle replication when the Okazaki-fragments are fully processed into a continuous strand.

We have previously reconstituted DNA replication with purified budding yeast proteins [5, 6] and here we have used this system to investigate replication into a nick. We introduced a sequence 1.7kb away from the origin which can be nicked on the leading or lagging strands by the Nb.BbvCI and Nt.BbvCI restriction nucleases, respectively. We first asked whether nascent DNA can be ligated efficiently to template DNA at a nick on the lagging strand template as outlined in Fig. 1A. We added ligase at different times after initiation to avoid ligation of the nick before the fork reaches it. We also omitted the lagging strand synthesis machinery (DNA polymerase δ (Pol δ), RFC, PCNA and Fen1) to prevent larger Okazaki fragments from confusing the analysis. Supplementary Figure 1A shows that a novel product larger than full-length plasmid (~9.4kb) was generated when ligase was added at 20 or 40min after initiation. This is the size predicted if the nascent leading strand was ligated to the nicked template. To verify that nascent strand-template hybrids can be generated, we replicated a linearised plasmid with a nick on the lagging strand (Supp. Fig. 1B,C). The replicated products were digested with the Dpn1 restriction enzyme. Dpn1 does not digest hemi-methylated DNA, but efficiently digests input plasmid DNA which is fully methylated in *E.coli*. Completely replicated products digested with DpnI should generate a full-length DNA strand in alkaline agarose gels (Supp. Fig. 1B), and this is what is seen when either unnicked template is replicated or ligase is added at time t=0 (Supp. Fig. 1C compare lanes 1-4 with 5-8 and 12,13 with 21,22). If the nascent strand is ligated to the template DNA, the full-length labelled product should be part DpnI-sensitive and part DpnI-resistant (Supp. Fig. 1B). Indeed, when ligase was added to the nicked template 20 or 40 min after initiation, the full-length products (Supp. Fig. 1C lanes 14-17) were lost after DpnI digestion (Supp. Fig. 1C lanes 23-26), and a novel band of ~2.3 kb was generated. Therefore, nascent DNA at a collapsed fork can be efficiently ligated to a nicked template.

To test whether a replication fork from the opposite direction can effectively re-replicate this ligated template, we replicated a circular plasmid nicked on the leading strand with the complete replication system including the Okazaki fragment synthesis and maturation machinery (Pol δ, RFC, PCNA and Fen1); ligase was added at either t=0 or t=20 min. When ligase was added at 20 min, in addition to full-length products, high MW rolling circle replication products were generated by 60 min (Fig. 1B lane 12). Taken together, these data show that nascent DNA can be ligated to the template DNA after fork collapse, and this can provide a substrate for localised re-replication.

Our results indicate that a collapsed fork can be a source of localized re-replication. We suggest that mechanisms must exist to regulate ligation of this nick so that it only happens in conjunction with replication restart. Perhaps poly ADP ribose polymerase (PARP), which binds tightly to nicks and recruits repair factors, plays a role in this regulation [7]. PARP is absent from yeast, so other mechanisms may also exist. Nicks can come from multiple sources including incomplete DNA repair and topoisomerase reactions. Structures like Fig. 1A ii can also be generated from stalled replication forks by structure-specific nucleases including Mus81 [8]. Consequently, mechanisms involved in regulating this ligation event are likely to be important for genome stability.

## Experimental Procedures

The experimental procedures are described in the supplemental information

## Author Contributions

E.J. executed the original experiments, E.J. and J.F.X.D. conceived the experiments, analysed the data and wrote the paper.

## Acknowledgements

We thank Max Douglas and Gideon Coster for plasmids and advice, and we thank the Fermentation Facility for growing yeast cultures. This work was supported by the Francis Crick Institute, which receives its core funding from Cancer Research UK (FC001066), the UK Medical Research Council (FC001066), and the Wellcome Trust (FC001066). This work was also supported by the Wenner-Gren Foundations (E.J) along with a Wellcome Trust Senior Investigator Award (106252/Z/14/Z) (J.F.X.D.) and a European Research Council Advanced Grant (669424-CHROMOREP) (J.F.X.D.).

