## Supplemental for "Can a nick promote partial genome re-replication?"

### Supplemental Material

### Experimental procedures

Over-expression of recombinant proteins

Cdc6, GINS, Mcm10, PCNA were over-expressed in bacteria as described previously[1-3]. BL21(DE3)-CodonPlus-RIL cells carrying the Sld2 over-expression plasmid were grown in LB with 100 µg/ml Amp and 37 µg/ml Chloramphenicol at 37 °C. At $OD_{600nm}$=0,5 were cells cooled to 16 °C and 0,2 mM IPTG was added to induce over-night expression.

ORC, Mcm2-7/Cdt1, DDK, S-CDK, Cdc45, Dpb11, Sld3/Sld7, Pol ε, RPA, Pol α, Ctf4, MrcI, Csm3/Tof1, TopoI, RFC, Pol δ, Fen1, Lig1 were over-expressed in yeast as described previously in [1-5].

Purification of proteins

ORC was purified as previously described in [1], with the exception that the peak fractions from the HiLoadSuperdex 16/600 were pooled and concentrated over an Amicon Ultra-15 100,000 NMWL concentrator instead of the previously described MonoQ column. ORC was stored in 25 mM HEPES-KOH (pH 7.6), 10 % glycerol, 200 mM KOAc, 0,05% NP-40-S, and 1 mM dithiothreitol.

Cdc6 was purified as previously described in [1], with the exception that Cdc6 in the last step was dialyzed against 25 mM HEPES-KOH (pH 7.6), 10 % glycerol, 100 mM KOAc, 10 mM $Mg(OAc)_2$, 0,02% NP-40-S and 1 mM dithiothreitol,

Mcm2-7/Cdt was purified as previously described in [5].

DDK was purified as previously described in [4] with the following modifications. An ATP-wash (25 mM HEPES-KOH (pH7.6), 10 % Glycerol, 400 mM NaCl, 0,02% NP-40-S, 1 mM dithiothreitol, 2 mM $CaCl_2$, 10 mM $Mg(OAc)_2$, and 1 mM ATP) was introduced when DDK was bound to the Calmodulin affinity resin. After washing with 20 bed volumes in the same buffer (without $Mg(OAc)_2$ and 1 mM ATP) was lambda phosphatase and 1 mM MnCl added to the column and left for 1 hour. Eluted DDK was concentrated over an Amicon Ultra-15 10,000 NMWL concentrator and passed over a Superdex200 equilibrated in 25 mM HEPES-KOH (pH 7.6), 10 % glycerol, 200 mM potassium glutamate, 0,02% NP-40-S, and 1 mM dithiothreitol.

S-CDK was purified as previously described in [2] with the following modifications. The TEV-protease was removed by passing the eluate from the calmodulin affinity resin over a Ni-NTA agarose column. The flow-through was concentrated over an Amicon Ultra-15 10,000 NMWL concentrator and passed over a Superdex200 column, equilibrated in 40 mM HEPES-KOH (pH 7.6), 10% glycerol, 300 mM KOAc, 0,02% NP-40-S, and 1 mM dithiothreitol.

Cdc45 was purified as previously described in [2] with the following modifications. The eluate from the anti-flag M2 affinity gel was concentrated over an Amicon Ultra-15 30,000 NMWL concentrator and passed over a Superdex200 column, equilibrated in 25 mM HEPES-KOH (pH 7.6), 10% glycerol, 300 mM KOAc, 1 mM Ethylenediaminetetraacetic acid (pH 8.0) (EDTA), and 1 mM dithiothreitol.

Dpb11 was purified as previously described in [2] with the following modifications. The eluate from the anti-flag M2 affinity gel was concentrated over an Amicon Ultra-15 3,000 NMWL concentrator and passed over a Superdex200 column, equilibrated in 25 mM HEPES-KOH (pH 7.6), 10% glycerol, 300 mM KCl, 1 mM EDTA, 0,02% NP-40-S, and 1 mM dithiothreitol.

GINS was purified as previously described in [2] with the following modifications. The lysate was incubated with Ni-NTA affinity resin in 50 mM HEPES-KOH (pH 7.6), 10 % glycerol, 200 mM KCl, 1 mM EDTA, 1 mM EGTA, 10 mM $Mg(OAc)_2$, 0,05% NP-40-S, 1 mM dithiothreitol, 0.5 mM AEBSF, 1 mM leupeptin, and 1 mM Pepstatin A. The resin was collected on a column, washed in the same buffer without protease inhibitors and with the addition of 10 mM Imidazol and 1 mM ATP. Bound proteins were eluted in the same buffer with 200 mM Imidazole. The eluate was pooled and dialyzed against 50 mM HEPES-KOH (pH 7.6), 10 % glycerol, 50 mM

KCl, 1 mM EDTA, 1 mM EGTA, 10 mM $Mg(OAc)_2$, 0,05% NP-40-S, and 1 mM dithiothreitol,. The sample was loaded on a 1 ml MonoQ column and eluted over a 30 CV gradient to 500 mM KCl. Peak fractions were concentrated and passed over a Superdex200 column, equilibrated in 25 mM HEPES-KOH (pH 7.6), 10 % glycerol, 200 mM KOAc, 1 mM EDTA, 0,02% NP-40-S, and 1 mM dithiothreitol.

Sld3/Sld7 was over-expressed and purified as described in [2].

Sld2 was over-expressed from plasmid pGC441 (a kind gift from Gideon Coster) in BL21(DE3)-CodonPlus-RIL cells. The lysate was incubated with Glutathione Sepharose in 25 mM HEPES-KOH (pH 7.6), 10% glycerol, 500 mM NaCl, 1 mM EDTA, 0.02% NP-40-S, 0.1% Tween20, 1 mM dithiothreitol, and the recommended concentration of a protease cocktail from cOmplete Mini, EDTA-free (Roche). After extensive washing, Sld2 was released from the affinity resin by incubation with PreScission protease. The eluate was pooled and diluted with an equal volume of buffer with no salt to reduce the salt concentration to 250 mM NaCl, and thereafter loaded onto a 1 mM HiTrap SP FF column equilibrated in 25 mM HEPES-KOH (pH 7.6), 10% glycerol, 250 mM NaCl, 1 mM EDTA, 0.02% NP-40-S, 0.1% Tween20, and 1 mM dithiothreitol. The peak fractions from a 10 CV step elution in the same buffer with 700 mM NaCl were pooled and dialyzed against 25 mM HEPES-KOH (pH 7.6), 40% glycerol, 700 mM KOAc, 1 mM EDTA, 0.02% NP-40-S, and 1 mM dithiothreitol.

Pol ε was purified as previously described in [2], but passed over a Superdex200 column, equilibrated in 25 mM HEPES-KOH (pH 7.6), 10% glycerol, 500 mM KOAc, and 1 mM dithiothreitol.

Mcm10 was purified as previously described in [2] with the following modifications. The lysate was incubated with anti-flag M2 affinity resin in 25 mM HEPES-KOH (pH 7.6), 10 % glycerol, 500 mM NaCl, 0,05% NP-40-S, 1 mM dithiothreitol, and the recommended concentration of a protease cocktail from cOmplete Mini, EDTA-free (Roche). The eluate from the anti-flag M2 affinity resin was pooled and 10 mM Imidazole was added, before incubation with Ni-NTA affinity resin. The eluate from the Ni-NTA affinity resin was concentrated over an Amicon Ultra-

15 30,000 NMWL concentrator and passed over a Superdex200 column, equilibrated in 25 mM HEPES-KOH (pH 7.6), 10 % glycerol, 200 mM NaCl, 1 mM EDTA, 0,05% NP-40-S, and 1 mM dithiothreitol.

RPA was purified as previously described in [2] with the following modifications. The lysate was incubated with calmodulin affinity resin in 25 mM Tris-HCl (pH 7.2), 10 % glycerol, 500 mM NaCl, 1 mM dithiothreitol, 0.5 mM AEBSF, 1 mM leupeptin, 1 mM Pepstatin A and 2 mM $CaCl_2$. An ATP wash was introduced in the same buffer (-protease inhibitors) with 10 mM $Mg(OAc)_2$ and 1 mM ATP. The eluate from the calmodulin affinity resin was diluted 2-fold to reduce the ionic strength to 250 mM NaCl, before loading onto a 1 ml HiTrap heparin column. Peak fractions from the 20 CV gradient were concentrated over an Amicon Ultra-4 10,000 NMWL concentrator and passed over a Superdex 200 column, equilibrated in 25 mM Tris-HCl (pH 7.2), 10 % Glycerol, 150 mM NaCl, , 1 mM EDTA and 1 mM dithiothreitol.

Pol $\alpha$ was purified as previously described in [2] with the following modification. An ATP-wash (25 mM Tris-HCl (pH 7.2), 10 % Glycerol, 400 mM NaCl, 0,01% NP-40-S, 1 mM dithiothreitol, 2 mM $CaCl_2$, 10 mM $Mg(OAc)_2$, and 1 mM ATP) was introduced when Pol $\alpha$ was bound to the Calmodulin affinity resin.

Ctf4 was purified as previously described in [2], but the peak fractions from the MonoQ column was concentrated over an Amicon Ultra-15 30,000 NMWL concentrator and passed over a Superdex200 column, equilibrated in 25 mM Tris-HCl (pH 7.2), 10 % Glycerol, 150 mM NaCl, 1 mM EDTA and 1 mM dithiothreitol. Peak fractions were pooled and concentrated over a Amicon Ultra-15 30,000 NMWL concentrator.

Mrc1 was purified as described in [3] with the following modification. An ATP-wash (25 mM Tris-HCl (pH 7.2), 10 % Glycerol, 400 mM NaCl, 1 mM EDTA, 0,02% NP-40-S, 1 mM dithiothreitol, 10 mM $Mg(OAc)_2$, and 1 mM ATP) was introduced when Mrc1 was bound to the anti-flag M2 affinity resin.

Csm3/Tof1 was purified as previously described in [3] with the following modifications. An ATP-wash (25 mM Tris-HCl (pH 7.2), 200 mM NaCl, 10 % Glycerol, 1 mM dithiothreitol, 2 mM $CaCl_2$, 10 mM $Mg(OAc)_2$, and 1 mM ATP) was introduced when Csm3/Tof1 was bound to the Calmodulin affinity resin. Next, the resin was resuspended in wash buffer and TEV was added to 100 $\mu$g/ml followed by incubation on ice for 2 hours. The eluate from the Calmodulin affinity resin was concentrated and applied to a Superdex 200 column, equilibrated in 25 mM Tris-HCl (pH 7.2), 150 mM NaCl, 10 % Glycerol, 0,02% NP-40-S, and 1 mM dithiothreitol.

TopoI was purified as previously described in [3] with the following modifications. In this simplified protocol, an ATP-wash (25 mM Tris-HCl (pH 7.2), 10 % Glycerol, 300 mM NaCl, 2 mM $CaCl_2$, 10 mM $Mg(OAc)_2$, 0,02% NP-40-S, 1 mM dithiothreitol, and 1 mM ATP) was introduced when TopoI was bound to the Calmodulin affinity resin. The eluate from the calmodulin beads was concentrated over a Amicon Ultra-15 10,000 NMWL concentrator and passed over a Superdex 200 column, equilibrated in 25 mM Tris-HCl (pH 7.2), 10 % Glycerol, 150 mM NaCl, 0,02% NP-40-S, and 1 mM dithiothreitol.

PCNA was purified as described in [3].

RFC was purified as previously described in [3] with the following modifications. An ATP-wash (25 mM HEPES-KOH (pH 7.6), 10 % Glycerol, 400 mM NaCl, 2 mM $CaCl_2$, 10 mM $Mg(OAc)_2$, 1 mM dithiothreitol, and 1 mM ATP) was introduced when RFC was bound to the Calmodulin affinity resin. The eluate from the Calmodulin affinity resin was loaded onto a 1 ml HiTrap SP column equilibrated in 25 mM HEPES-KOH (pH 7.6), 10 % Glycerol, 200 mM NaCl, and 1 mM dithiothreitol. Proteins were eluted with a 30 ml gradient to 25 mM HEPES-KOH (pH 7.6), 10 % Glycerol, 1 M NaCl, and 1 mM dithiothreitol. The eluate from the HiTrap SP column was concentrated over a Amicon Ultra-15 30,000 NMWL concentrator and passed over a Superdex200 column, equilibrated in 25 mM HEPES-KOH (pH 7.6), 10 % Glycerol, 150 mM NaCl, and 1 mM dithiothreitol.

Pol $\delta$ was purified as previously described in [3] with the following modifications. An ATP-wash (25 mM Tris-HCl (pH 7.2), 10 % Glycerol, 400 mM NaCl, 2 mM $CaCl_2$, 0,02% NP-40-S, 1 mM dithiothreitol, 10 mM $Mg(OAc)_2$, and 1 mM ATP) was introduced when Pol δ was bound to the Calmodulin affinity resin. The eluate from the Calmodulin affinity resin was loaded onto a 1 ml HiTrap Heparin column equilibrated in 25 mM Tris-HCl (pH 7.2), 10 % Glycerol, 200 mM NaCl, 1 mM EDTA, and 1 mM dithiothreitol. Proteins were eluted with a 15 ml gradient to 1 M NaCl in the same buffer. The eluate from the HiTrap heparin column was concentrated over a Amicon Ultra-15 30,000 NMWL concentrator and passed over a Superdex 200 column, equilibrated in 25 mM Tris-HCl (pH 7.2), 10 % Glycerol, 150 mM NaCl, 0,02% NP-40-S, and 1 mM dithiothreitol.

Fen1 was purified as previously described in [6], with the exception that Fen1 as a last step was dialyzed against 25 mM Tris-HCl (pH 7.2), 10 % glycerol, 100 mM NaCl, 0,02% NP-40-S, and 1 mM dithiothreitol.

Lig1 was purified as previously described in [6], with the exception that Lig1 as a last step was dialyzed against 50 mM Tris-HCl (pH 7.5), 10 % glycerol, 100 mM NaCl, 1 mM EDTA and 1 mM dithiothreitol.

The template for the in vitro replication assay

The plasmid used as a template, CEJ5, in the in vitro replication assay was derived from plasmid MD154 (a kind gift from Max Douglas). A linker with a unique recognition site for Nt.BbvCI and Nb.BbvCI was introduced about 1,7 kb from the eukaryotic origin of replication. Nt.BbvCI or Nb.BbvCI (New England Biolabs) was used to create a single nick in the plasmid, positioned on either the leading or lagging strand template for an approaching replication fork. A unique restriction site, ScaI on the opposite side of the origin allowed the plasmid to be linearized when indicated.

Replication assays

In vitro replication assays were performed as previously described [2, 3] but with the following modifications. MCM loading was carried out by incubating either circular or linearized 4 nM plasmid DNA (CEJ5) with 8 nM ORC, 15 nM Cdc6, 22,5 nM Mcm2-7/Cdt1, 5 mM ATP, 25 mM HEPES-NaOH (pH 7,6), 100 mM potassium glutamate, 10 mM $Mg(Oac)_2$, 2 mM dithiothreitol, and 0,02% NP-40-S at 30 °C for 10 minutes. Next, the reaction was supplemented with 20 nM Dbf4-dependent kinase (DDK) and incubated for an additional 5 minutes at 30 °C. The loading reaction was diluted two-fold upon the initiation of DNA replication when the following reagents were added (final concentrations): 5 nM S-CDK, 7,5 nM Dpb11, 15 nM GINS, 120 nM Cdc45, 15 nM Pol $\varepsilon$, 7,5 nM Mcm10, 5 nM Ctf4, 50 nM RPA, 15 nM Csm3/Tof1, 20 nM Mrc1, 5 nM TopoI, 80 nM Pol $\alpha$, 7,5 nM Sld3/Sld7, 9 nM Sld2, 200 $\mu$M CTP, 200 $\mu$M GTP, 200 $\mu$M UTP, 80 $\mu$M dATP, 80 $\mu$M dGTP, 80 $\mu$M dTTP, 80 $\mu$M dCTP and 1 $\mu$Ci [$\alpha$-$^{32}$P]-dCTP. The final buffer condition, including salt and glycerol supplemented by the added proteins was: 30 mM HEPES-NaOH (pH 7,6), 16 mM $Mg(Oac)_2$, 3 mM dithiothreitol, 0,02% NP-40-S, 11 mM KCl, 48 mM KAc, 250 mM potassium glutamate, and <2% glycerol. 20 nM ligase (Cdc9) was added as indicated and in addition 20 nM RFC, 100 nM PCNA, 5 nM Pol $\delta$, 20 nM Fen1 and increased the final KCl concentration to 60 mM when processing of Okazaki-fragments was required. Reactions were incubated at 30 °C for up to 60 minutes after which they were quenched by the addition of 65 mM EDTA before removal of unincorporated nucleotides using Illustra G-50 columns (GE Healthcare). The products were separated on a 1% alkaline agarose gel, run for 16 hours at 30 V as described earlier [3]. The alkaline agarose gels were fixed by incubation for 2x20 minutes in 5% trichloroacetic acid solution at room temperature before drying on 3MM chromatography paper. Gels were autoradiographed using Amersham Hyperfilm MP (GE Healthcare)

### Supplementary figure legends

#### *Supplementary Figure 1*

Supplementary Figure 1. (A) Probing whether nascent DNA is ligated to template DNA on a circular substrate when the first fork runs into a nick on the lagging strand template. The Okazaki-fragment machinery (Pol δ, RFC, PCNA and Fen1) was omitted. Ligase 1 was added 20 or 40 minutes after the initiation of DNA replication, giving the fork time to reach the nick before ligase was added. The longest product (1) is the result of a ligation event between the nascent DNA and template DNA. (B) The scheme illustrates possible outcomes when DpnI is incubated with replication products. DpnI cuts DNA purified from E.coli but not in vitro replicated hemi-methylated DNA. (C) Probing whether nascent DNA is ligated to template DNA on a linearized substrate when the fork enocunters a nick on the lagging strand template. The Okazaki-fragment machinery (Pol δ, RFC, PCNA and Fen1) was omitted. Ligase 1 was added 20 or 40 minutes after the initiation of DNA replication, giving the fork time to reach the nick before ligase was added. Replication products in lanes 5-7 and 18-26 were incubated with DpnI.

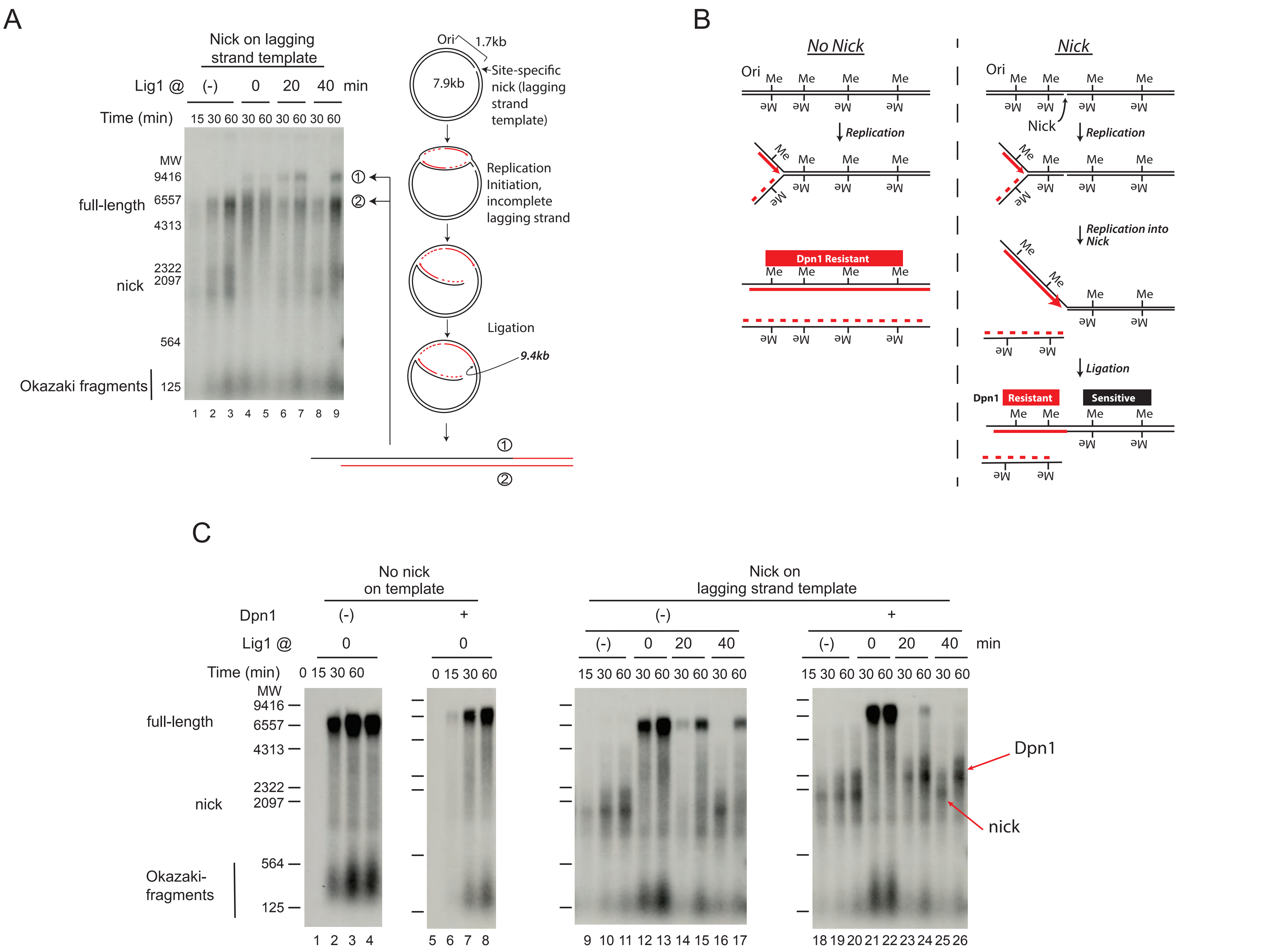
A
Nick on lagging strand template
Lig1 @ (-) 0 20 40 min
Time (min) 15 30 60 30 60 30 60 30 60
MW
9416
6557
4313
2322
2097
564
125
full-length
nick
Okazaki fragments
1 2 3 4 5 6 7 8 9
Ori
1.7kb
7.9kb
Site-specific nick (lagging strand template)
Replication Initiation, incomplete lagging strand
Ligation
9.4kb
B
No Nick
Nick
Ori
Me
Replication
Replication into Nick
Dpn1 Resistant
Ligation
Dpn1 Resistant Sensitive
C
No nick on template
Nick on lagging strand template
Dpn1 (-) +
Lig1 @ 0 0
Time (min) 0 15 30 60
(-) 0 20 40 min
15 30 60 30 60 30 60 30 60
full-length
nick
Okazaki-fragments
Dpn1
nick
